# GeomeTRe: accurate calculation of geometrical descriptors of tandem repeat proteins

**DOI:** 10.1101/2025.04.24.650440

**Authors:** Zarifa Osmanli, Elisa Ferrero, Alexander Miguel Monzon, Silvio C E Tosatto, Damiano Piovesan

## Abstract

Structured tandem repeat proteins (STRPs) are characterized by preserved structural motifs arranged in a modular way. The structural and functional diversity of STRPs makes them particularly important for studying evolution, novel structure and function relationships, and ultimately for designing new synthetic proteins with specific functions. One crucial aspect of their classification is the estimation of geometrical parameters, which can provide better insight into their properties and the relationship between the spatial arrangement of repeated units and protein function. Calculating geometric descriptors for STRPs is challenging because naturally occurring repeats are not perfect and often contain insertions and deletions. Existing tools for predicting structural symmetry work well on simple cases but often fail for most natural proteins. Here we present GeomeTRe, an algorithm that calculates geometrical descriptors such as curvature (yaw), twist (roll), and pitch for a protein structure with known repeat unit positions. The algorithm simulates the movement of consecutive units, identifies rotational axes, and calculates the corresponding Tait-Bryan angles. The parameters of GeomeTRe can enhance STRP annotation and classification by identifying variations in geometric arrangements among different functional groups. The package is fast and suitable for processing large protein structure datasets when repeat region information (e.g., from RepeatsDB) is available.

GeomeTRe is available as a Python package; source code and documentation can be found at https://github.com/BioComputingUP/GeomeTRe

## 1 Introduction

Tandem repeat proteins (TRPs) are prevalent throughout the tree of life and perform a broad range of functions (Marcotte et al. 1999, Kajava and Tosatto 2018). Structured tandem repeat proteins (STRPs) are a specific group characterized by conserved structural motifs in repetitive regions that do not necessarily share sequence similarity (Monzon et al. 2023). According to Kajava’s classification, STRPs can be grouped into five classes based on their shape and length (Kajava 2011). STRPs have distinctive structural and functional features, making them biologically significant subjects of study (Donagh et al. 2024, Monzon et al. 2023, Arrias et al. 2024).

A deep understanding of repeat proteins’ structural characteristics is important for uncovering biological insights and for applications in *de novo* protein design (Vrancken et al. 2020). In protein design, STRPs - especially alpha-helical repeats - are frequently used as starting scaffolds due to their relatively simple structure (Brunette et al. 2015, Doyle et al. 2015). Studies in alpha-helical protein design suggest that STRPs can serve as scaffolds for biomaterials in nanotechnology and biomedical applications (Parmeggiani 2015). For example, Park and colleagues controlled the curvature of repeat units to achieve a desired protein design (Park et al. 2015).In protein engineering, STRPs are valuable tools for biomedical applications due to their simplified sequence–structure relationships (Parmeggiani and Huang, 2017).

The RepeatsDB classification, which extends Kajava’s, subdivides solenoid STRPs into alpha, beta, and alpha–beta topologies (Clementel et al. 2025). Solenoids can be described as a continuous superhelix where the superhelix axis defines its “curvature” (Kobe and Kajava 2000). In this context, curvature results from the rotation of consecutive repeat units relative to each other, and is not necessarily aligned with the helical axis. Curvature can vary from large to small and serves as an additional parameter for STRP classification (Kobe and Kajava 2000). A recent review summarizes the classification of various alpha-solenoid subfolds based on curvature (Arrias et al. 2024). Twist is another geometrical property, defined as the coiling angle between consecutive units (Kobe and Kajava 2000). Twist can be right-handed (clockwise) or left-handed (anticlockwise). Most solenoid STRPs have an energetic preference for right-handed twists (Kobe and Kajava 2000), while others are left-handed, while others are left-handed (e.g., the LRR protein LegL7) (Batkhishig et al. 2021).

For solenoids, properties such as curvature, twist, and handedness have mostly been described using a superhelical model. However, existing methods do not fully cover the structural diversity of STRPs. Tools like SymD detect internal symmetry by aligning numerous fragments and applying circular permutations to find high-scoring matches, but they do not explicitly provide curvature, handedness, or twist values (Kim et al. 2010).

Similarly, CE-Symm predicts internal symmetry in elongated and closed repeat structures but cannot directly measure curvature or twist (Bliven et al. 2019).

Notably, Brunette and colleagues designed solenoids with controlled curvature by optimizing some global geometrical parameters defined by Kobe and Kajava (Kobe and Kajava 2000, Brunette et al. 2015). Another approach, HELFIT, uses a total least squares method to fit helical structures and defines features such as the helix axis, radius, pitch, handedness, and regularity (Enkhbayar et al. 2008).

Building on HELFIT, the recently proposed RepeatParam algorithm uses a helix-on-helix model to capture both the global architecture and the local superhelical structure of repeats (Pretorius and Murray, 2024). While effective for helical repeats, these approaches may not perform well on non-helical repeat structures.

Given these limitations, we developed GeomeTRe to characterize the geometrical properties of both open and closed-repeat proteins. The algorithm employs a circular fitting approach that provides a more flexible and accurate framework. The software is open-source and distributed as a Python package.

## 2 Materials and Methods

GeomeTRe calculates geometrical properties of tandem repeat proteins. It requires a protein structure and the start and end positions of each repeat unit (with optional insertion positions) as input. If insertion positions are provided, those segments are excluded to improve accuracy. For most known STRPs, repeat unit and insertion coordinates are available from the manually curated RepeatsDB database (Clementel et al. 2025) (Supp. Table S1). The algorithm computes the three Tait-Bryan angles - ***yaw, pitch***, and ***roll*** - (Humphreys 1912, Allgeuer and Behnke, 2015) by simulating an airplane traversing the protein from its N-terminus to C-terminus. In this analogy, the airplane points to the centroid of the next repeat unit, and the angles correspond to the maneuvers required to move from one unit to the next. The strategy for defining the rotation axes used to calculate these angles is described below. The algorithm also determines handedness, defined by the roll direction of movement (clockwise/right-handed or anticlockwise/left-handed), and the sign of the pitch (positive for upward, negative for downward movement).

Curvature and twist have historically been used to describe STRP structures (Kobe and Kajava 2000). In our airplane analogy, these correspond to yaw and roll; however, an unambiguous description of the motion also requires calculating the pitch angle. This analogy between aircraft motion and the relative orientation of protein repeat units is illustrated in Figure 1A.

**Figure 1.**
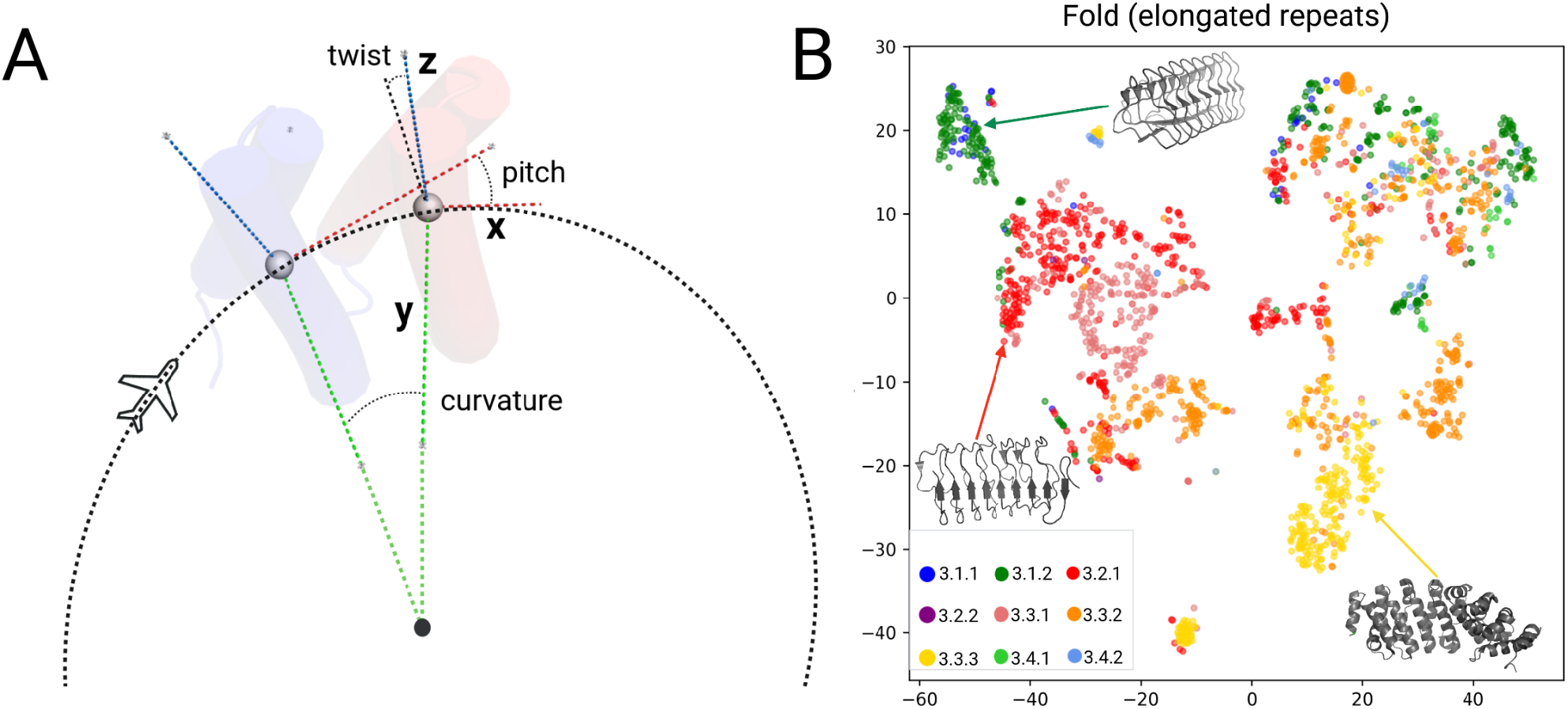
GeomeTRe parameter definitions and their distribution in elongated STRPs. **(A)** The algorithm fits a circle through the geocenters of the repeat units to identify the rotational axes. The twist axis (x, red) is tangent to the fitted circle. The pitch axis (y, green) points toward the circle’s center. The curvature axis (z, blue) is orthogonal to the other two axes. Angles are calculated by superimposing two consecutive units after these axes are defined. **B)** t-SNE clustering of class 3 elongated STRPs based on GeomeTRe output parameters (mean and standard deviation of curvature, twist, and pitch). Colors correspond to fold classifications in RepeatsDB (3.1.1 - right-handed beta solenoid; 3.1.2 - left-handed beta solenoid; 3.2.1 - high curvature alpha/beta solenoid; 3.2.2 - low curvature alpha/beta solenoid; 3.3.1 - low curvature alpha solenoid; 3.3.2 - high curvature alpha solenoid; 3.3.3 - corkscrew alpha solenoid; 3.4.1 - multimeric beta-hairpin; 3.4.2 - monomeric beta-hairpin).

Identifying and labeling rotation axes in proteins is not straightforward. Natural protein structures are irregular, and repeat units can lie on shapes for which it is difficult to define a dominant axis. Given the high degeneracy and structural variability among units in natural repeats, we divided the problem and made certain simplifying assumptions.

### Curvature (yaw)

In GeomeTRe, the roll axis (direction of motion) corresponds to the vector connecting the centroids of two consecutive units. To identify the other two axes, we define a reference plane by fitting a circle through the repeat units. This circle defines the reference plane and is also used to calculate the structure’s curvature.

To account for variable protein shapes and ensure numerical stability, the algorithm uses a “widest circle” approach with a sliding window of six units. The plane is determined by the first two principal components of the centroids (geocenters) of the units in the window (these correspond to the roll/twist and pitch axes). All Cα atoms of the window’s units are projected onto this plane, and the largest circle that fits within the innermost and outermost projected atoms of each unit is identified (Figure 1A). Finally, for each unit, the largest such circle among all overlapping windows is selected and its center is computed. Curvature is defined as the angle at the circle’s center subtended by the centroids of two consecutive units, and is computed for each unit. Using a sliding window allows analysis of superhelical structures without accumulating large errors.

### Twist (roll), pitch and handedness

Twist and pitch angles are obtained by structurally aligning each repeat unit to its predecessor using TM-align (Zhang and Skolnick, 2005). To ensure accuracy, we first rotate both units to a reference orientation defined by their pitch and twist axes. For each unit, the pitch axis is defined as the vector from the center of the fitted circle to the unit’s centroid (i.e., the circle’s radius). The twist axis is computed by orthogonalizing the vector connecting two consecutive centroids with respect to the pitch axis, yielding the tangent to the circle at that unit. Prior to alignment, we rotate each unit by the cross-product of its twist and pitch axes to standardize their orientation.

After alignment, the resulting rotation is decomposed into twist (roll), pitch, and curvature (yaw) components. We discard the yaw component from this alignment, since curvature is already determined by the circle-fitting method. Twist handedness is defined as positive for clockwise rotations (negative for anticlockwise), and pitch handedness is positive when a unit moves upward (negative for downward).

### Implementation

GeomeTRe is available as a Python package and can be used as a command-line tool or as a library. It relies on standard Python libraries (Numpy, Pandas, SciPy, scikit-learn, scikit-image, BioPython) and the *tmtools* package (which wraps TM-align).GeomeTRe also provides an option to visualize the rotational axes by launching a PyMOL session.

The program takes a protein structure and the start–end positions of its repeat units (and any insertions) as input. Repeat annotations for most known STRPs can be retrieved from RepeatsDB. The output is the set of geometrical parameters for each unit, returned as a Pandas DataFrame (which can be printed to standard output or saved to a CSV file).

GeomeTRe can also run in batch mode to process multiple PDB files in parallel. The algorithm is fast, taking less than a second per structure. The package is open-source under a GPLv3 license and is available at https://github.com/BioComputingUP/GeomeTRe.

## 3 Results and Discussion

The GeomeTRe algorithm accurately determines the curvature of elongated and closed repeats, as well as challenging cases like linear structures with very low curvature. We analyzed the distribution of GeomeTRe parameters for all experimentally derived STRPs in RepeatsDB (Supp. Figs S1-S5). Extended structures (class 3) vary in their number of repeat units, whereas closed STRPs (class 4) are constrained by their shape. This is reflected in the variability of geometrical parameters, especially curvature. For example, closed topologies like TIM barrels (class 4.1) and propellers (class 4.4) always have curvature above 0.5 radians, whereas elongated repeats like beta-solenoids (class 3.1) are rod-like with limited flexibility and exhibit very low curvature (Supp. Fig. S1). For instance, the ice-binding protein (PDB 6eio), which helps protect bacteria from freezing (Mangiagalli et al. 2018), has a beta-solenoid structure with minimal curvature.

For most topologies, pitch, twist, and curvature are positively correlated (Supp. Figs S6–S8). The notable exception is the trefoil topology (class 4.3), where curvature is inversely correlated with both pitch and twist (Supp. Figs S7–S8). Overall, GeomeTRe parameters exhibit only weak inter-correlations, suggesting they capture distinct aspects of STRP architecture. To explore this, we performed t-SNE clustering on class 3 elongated repeats using these parameters (Figure 1B). The resulting clusters align not only with the broad topology classes but also with specific fold subtypes.

We also observed that the mean of each parameter is positively correlated with its standard deviation across all topologies, except for class 4.3. The outlier behavior of topology 4.3 can be explained by the presence of two distinct fold populations in this class, which differ substantially in how their repeat units are organized.

### Natural versus designed STRPs

In a “perfect repeat,” the rotations between consecutive units are nearly identical for all pairs. In natural proteins, however, these rotations can vary significantly depending on the degree of structural variability between units.

We compared the geometrical parameters of “natural” STRPs versus designed repeat proteins. We assembled a dataset of 8,622 natural STRP structures (15,180 repeat regions) from RepeatsDB v4 (https://repeatsdb.org/) and 76 designed repeat structures (132 repeat regions) generated using Rosetta (Brunette et al. 2015.). We then compared their parameter distributions (Supp. Figs S12–S13). Table 1 provides a direct comparison of GeomeTRe and Rosetta script estimates for twist, pitch, and curvature on a representative set of designed helical repeat (DHR) proteins.

**Table 1.**
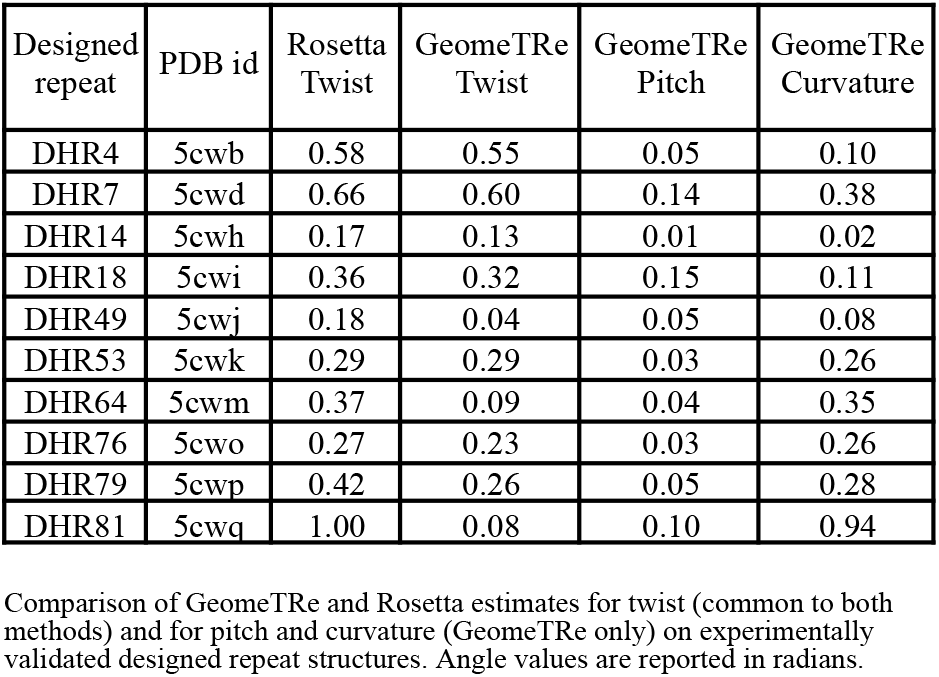
Comparison of GeomeTRe and a Rosetta on designed helical repeats.

Natural proteins show a moderate positive correlation between mean twist and mean pitch, whereas designed STRPs exhibit a much stronger correlation between these parameters (Supp. Fig. S13). The distribution of twist and curvature also differs: designed proteins cover a much narrower range (Supp. Fig. S12). Natural STRPs display a broad range of curvature, with mean values spanning ∼0 to 2 radians (Supp. Fig. S12). In natural elongated STRPs, strong parameter correlations are observed only at the topology level (primarily between twist and pitch). In contrast, designed STRPs show consistently high correlations among parameters (Supp. Fig. S13). These pronounced correlations in designed proteins may indicate that structural constraints were optimized during design for specific functional or stability requirements.

All input data and results from this study are available at https://github.com/BioComputingUP/GeomeTRe_results.

## 4 Conclusions

We have presented GeomeTRe, a Python package for fast and accurate calculation of the main geometrical properties of tandem repeat proteins using their structures and repeat annotations from RepeatsDB. The package calculates curvature, twist, pitch, and handedness, providing valuable insights for STRP classification. Our circular fitting method outperforms a superhelical fitting approach, particularly for β-solenoids and closed repeats, which have low curvature and a more planar organization. The geometric parameters computed by GeomeTRe provide a quantitative description of repeat protein structures, enabling more refined structural classifications and potential refinement of the current RepeatsDB classification.

