## Supplementary Data for "GeomeTRe: accurate calculation of geometrical descriptors of tandem repeat proteins"

### Table of contents

**Table of contents** 1
**1. RepeatsDB classification** 2
**2. Parameters distributions** 3
**3. Parameters correlations** 8
**4. Mean and standard deviation (std) correlations** 11
**5. Comparison between natural and designed STRPs** 14

### 1. RepeatsDB classification

**Table S1**. Classification of elongated and closed structured tandem repeat proteins (STRPs) in RepeatsDB (repeatsdb.org).

| Classification id | Classification name in topology level |
|---|---|
| 3.1 | Beta-solenoid |
| 3.2 | Alpha/beta solenoid |
| 3.3 | Alpha-solenoid |
| 3.4 | Beta hairpins |
| 4.1 | TIM-barrel |
| 4.2 | Beta-barrel/beta hairpins |
| 4.3 | Trefoil |
| 4.4 | Propeller |
| 4.5 | Alpha/beta prism |
| 4.6 | Alpha-barrel |
| 4.7 | Alpha/beta barrel |
| 4.8 | Aligned prism |

#### 2. Parameters distributions

Topology 4.7 is excluded in Figure S1-S3 due to the small number of representative STRPs available in RepeatsDB.

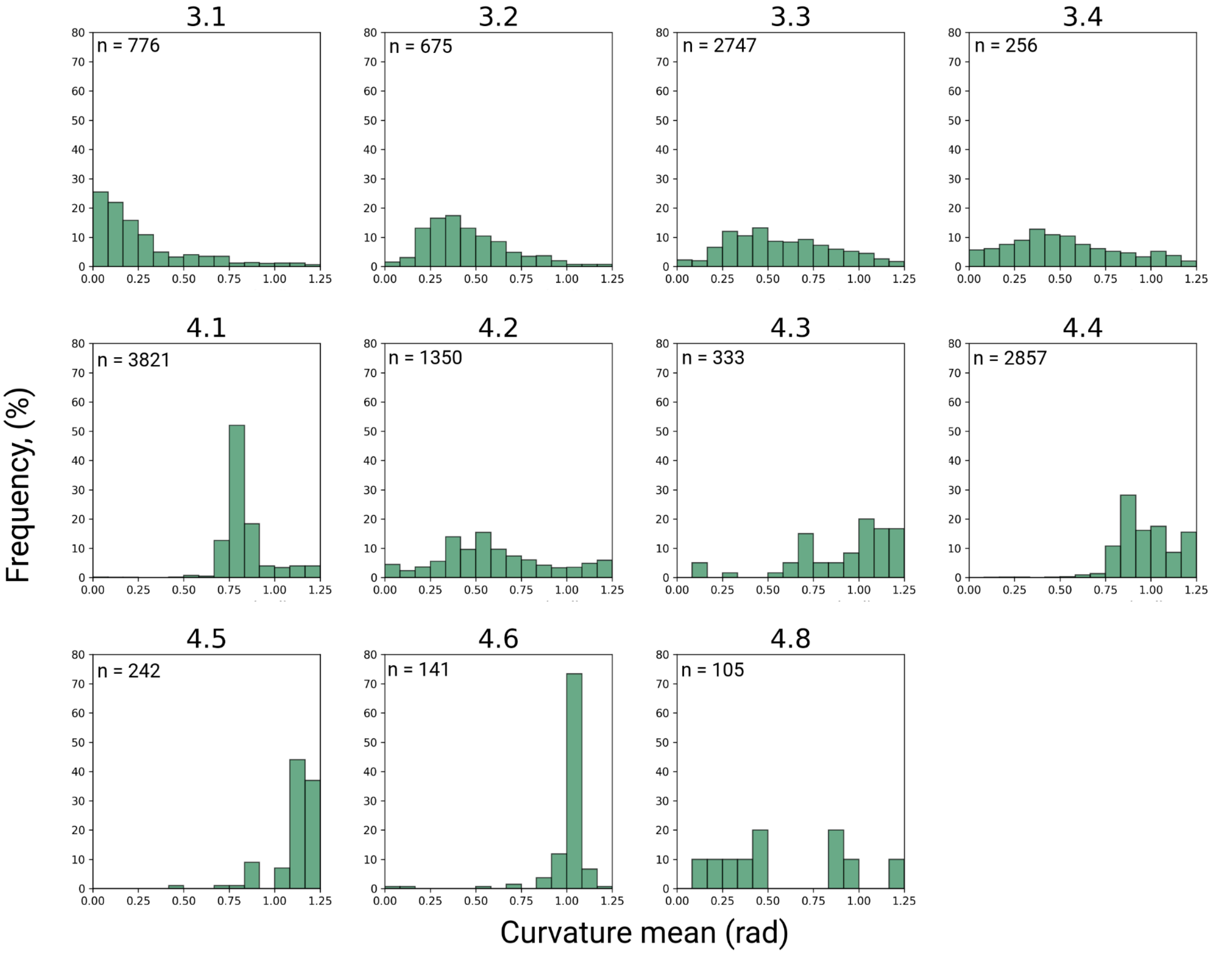


**Figure S1**. Frequency distribution of mean of curvature parameter across topologies of elongated (class 3) and closed (class 4) STRPs.

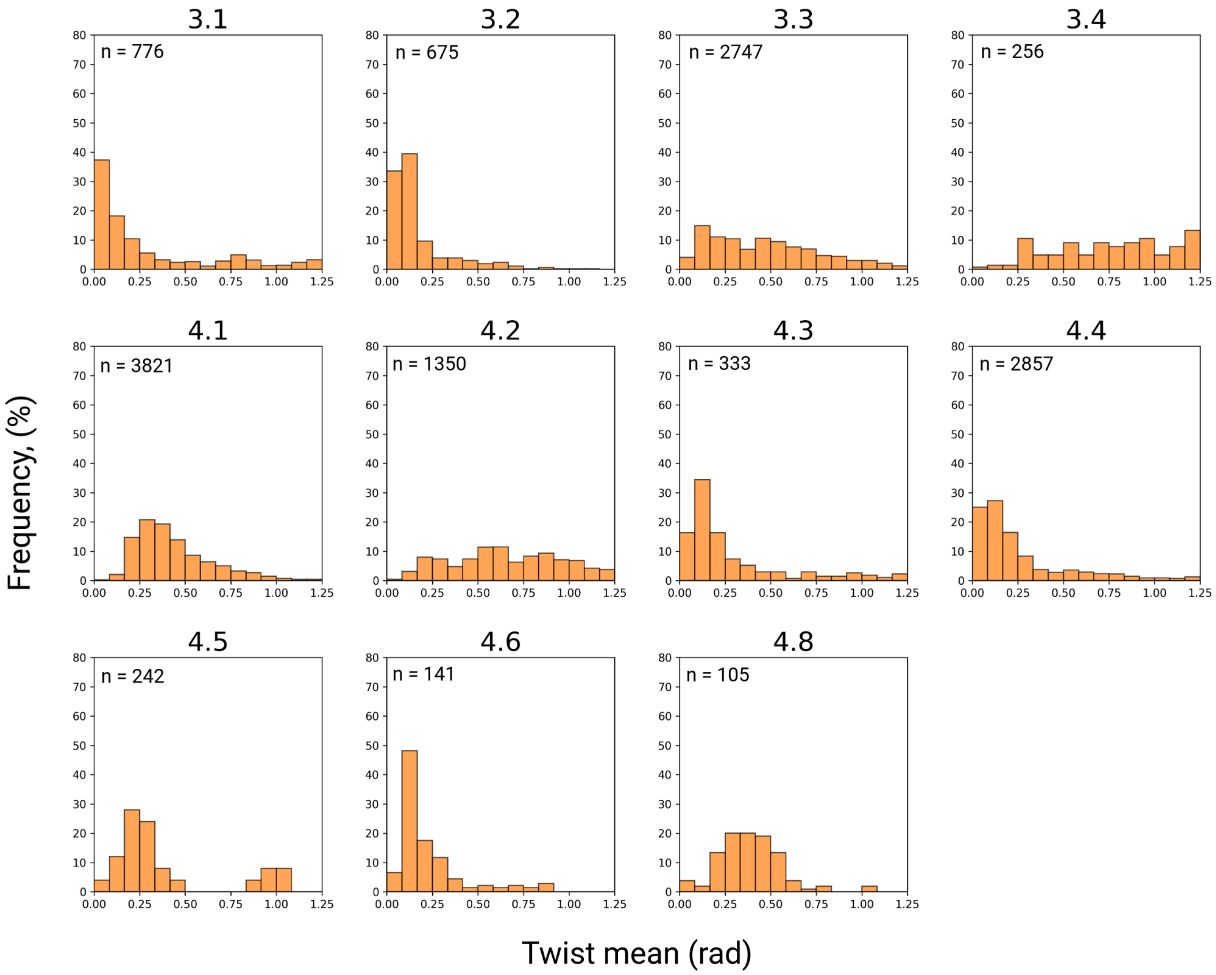


**Figure S2**. Frequency distribution of mean of twist parameter across topologies of elongated (class 3) and closed (class 4) STRPs.

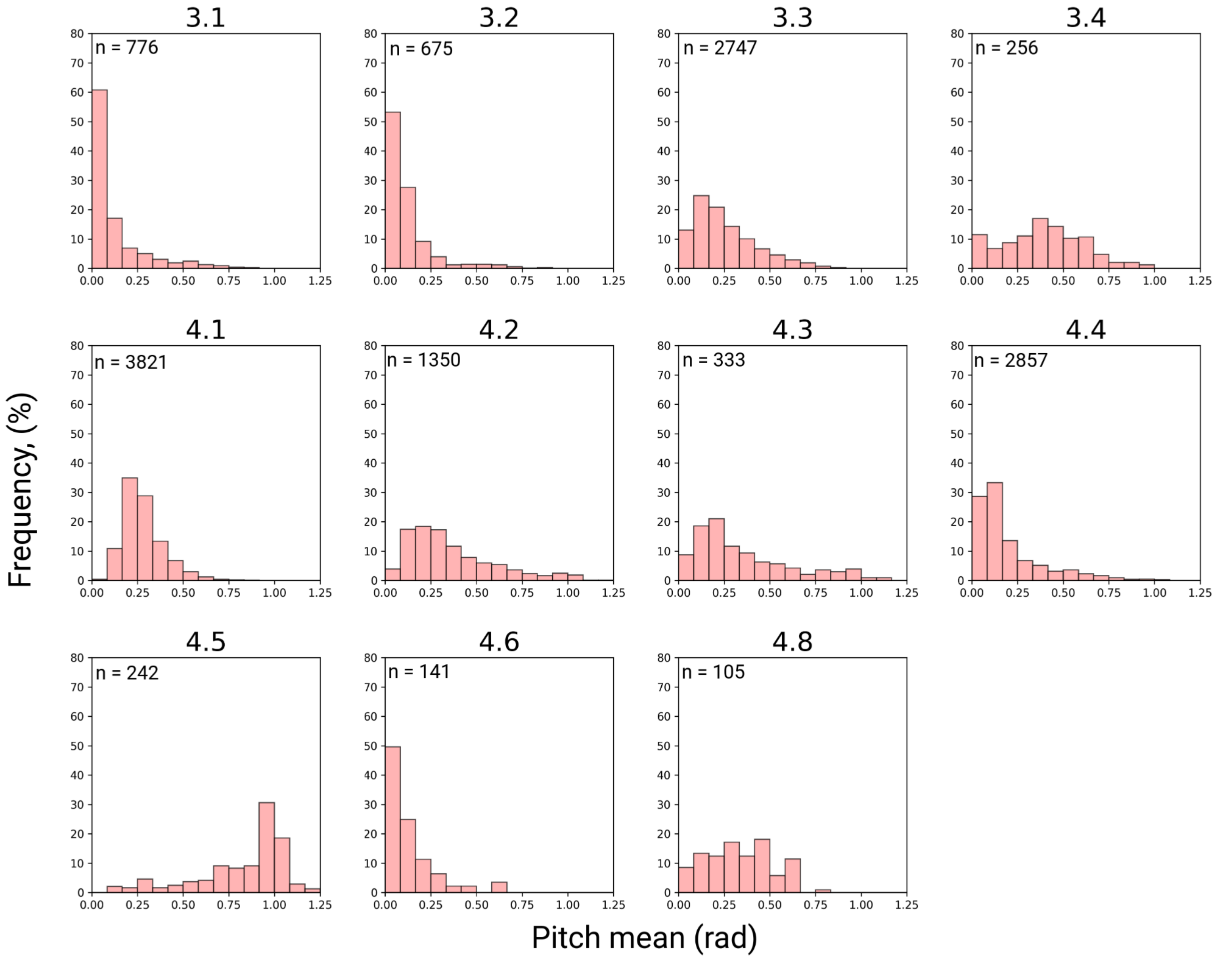


**Figure S3**. Frequency distribution of mean of pitch parameter across topologies of elongated (class 3) and closed (class 4) STRPs.

The handedness of the protein structure can be either right handed (+1) or left handed (-1). For some STRPs the handedness changes direction along the region and resulting average values are continuous. For the handedness statistics in order to exclude structures which are too linear or that change direction, we considered only STRP regions with both an average higher than 0.1 and a standard deviation lower than 0.1. Structures with less than 6 units are excluded. None of the structures in topologies 4.7 and 4.8 passed those filtering criteria and therefore are not shown.

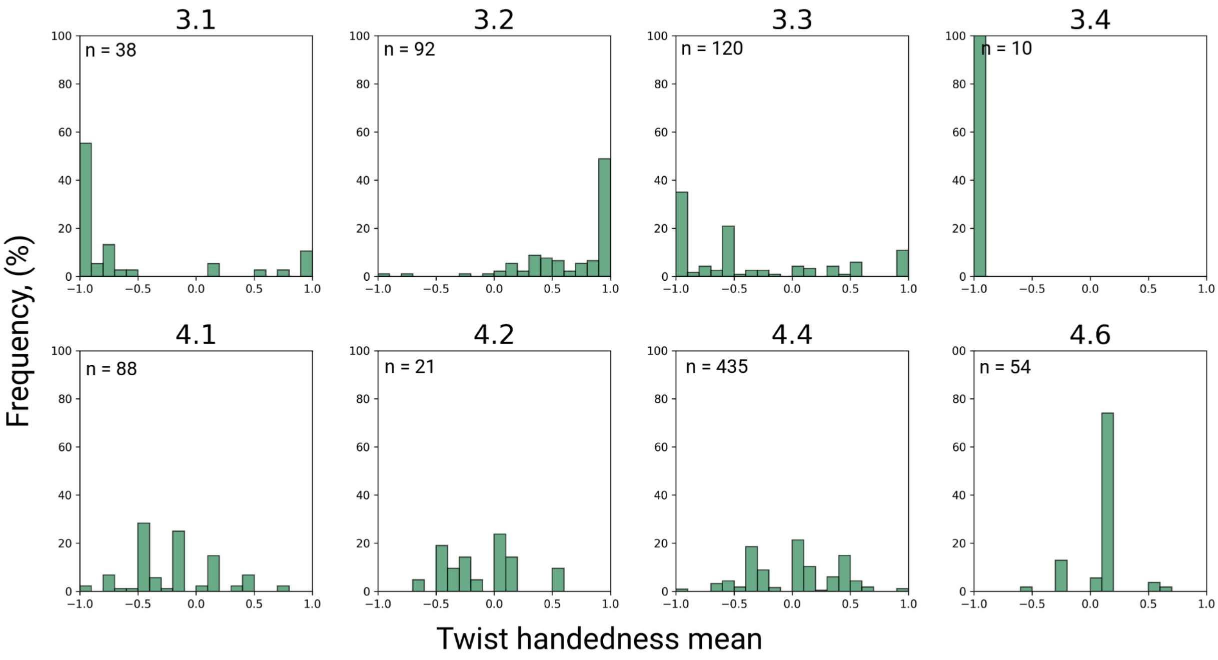


**Figure S4**. Frequency distribution of mean of twist handedness among topologies of elongated (class 3) and closed (class 4) STRPs.

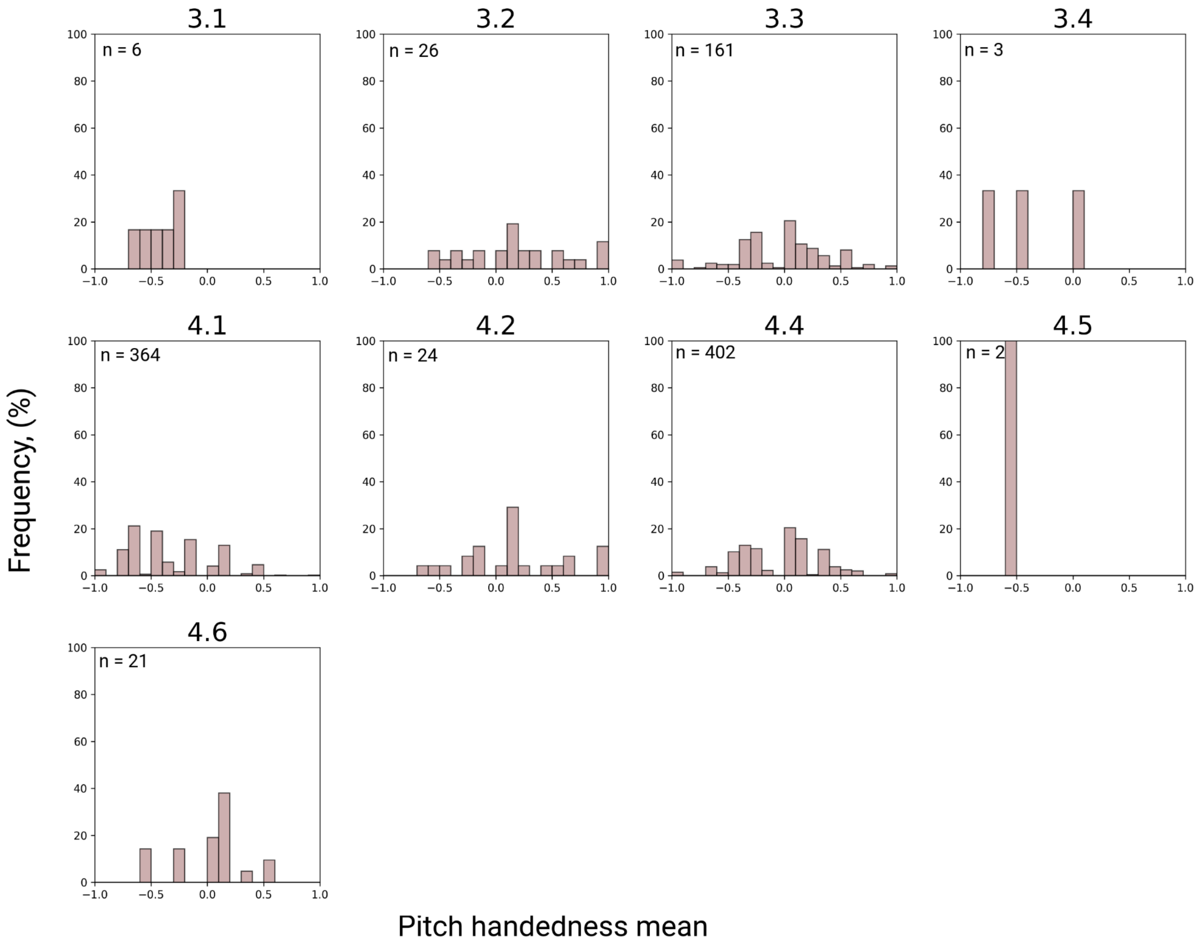


**Figure S5**. Frequency distribution of mean of pitch handedness among topologies of elongated (class 3) and closed (class 4) STRPs.

#### 3. Parameters correlations

P-value significance of each topology is specified with asterisks (*). $p < 0.001$ is "***", $p < 0.01$ is "**", $p < 0.05$ is "*" $p >= 0.05$ is "ns" (not significant).

Topology 4.7 is excluded due to the small number of available STRPs in RepeatsDB. The number of repeat regions considered in each topology (subplot) is indicated with *n*.

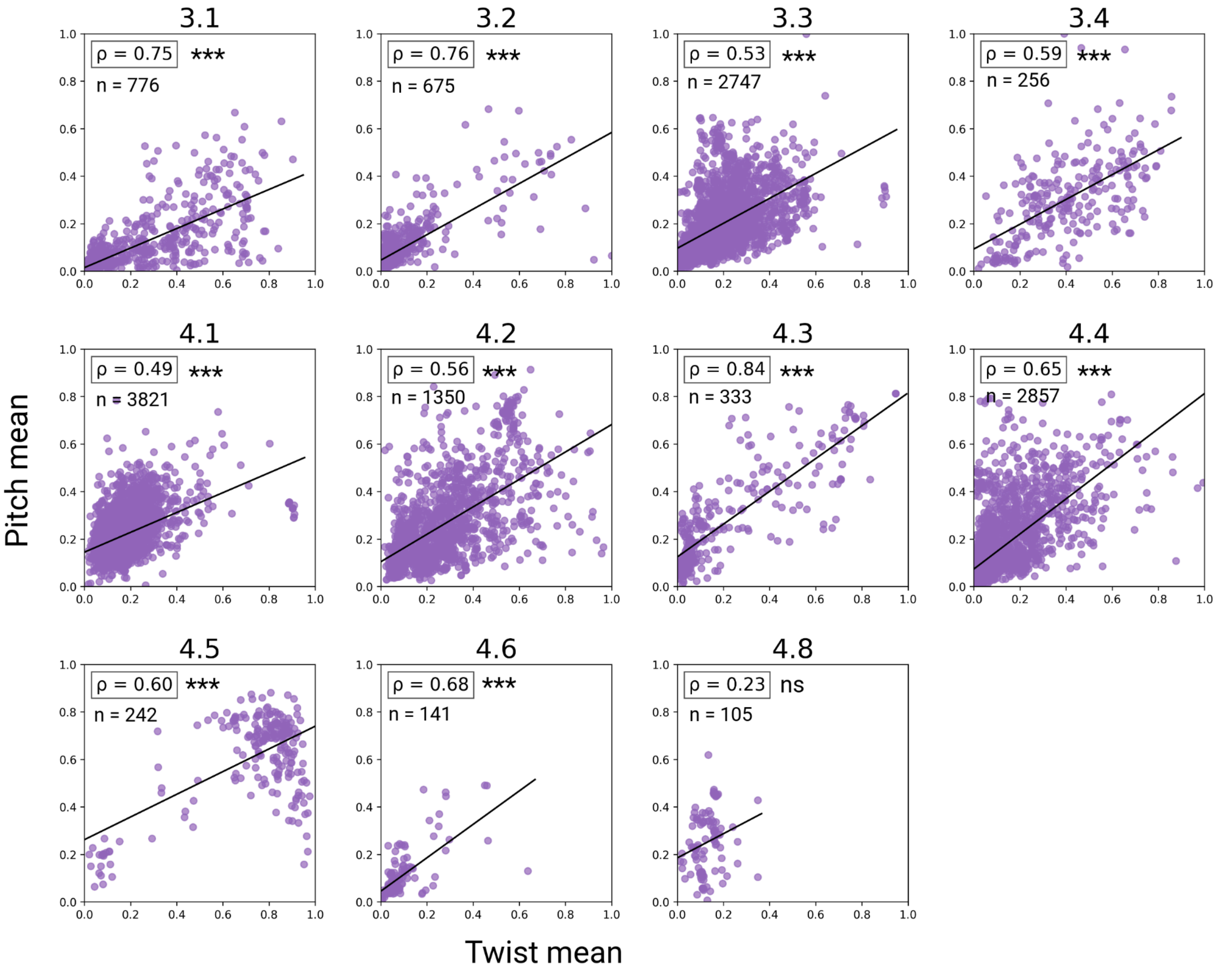


Figure S6. Correlation of mean of pitch and twist parameters for elongated (class 3) and closed (class 4) STRPs at the topology level.

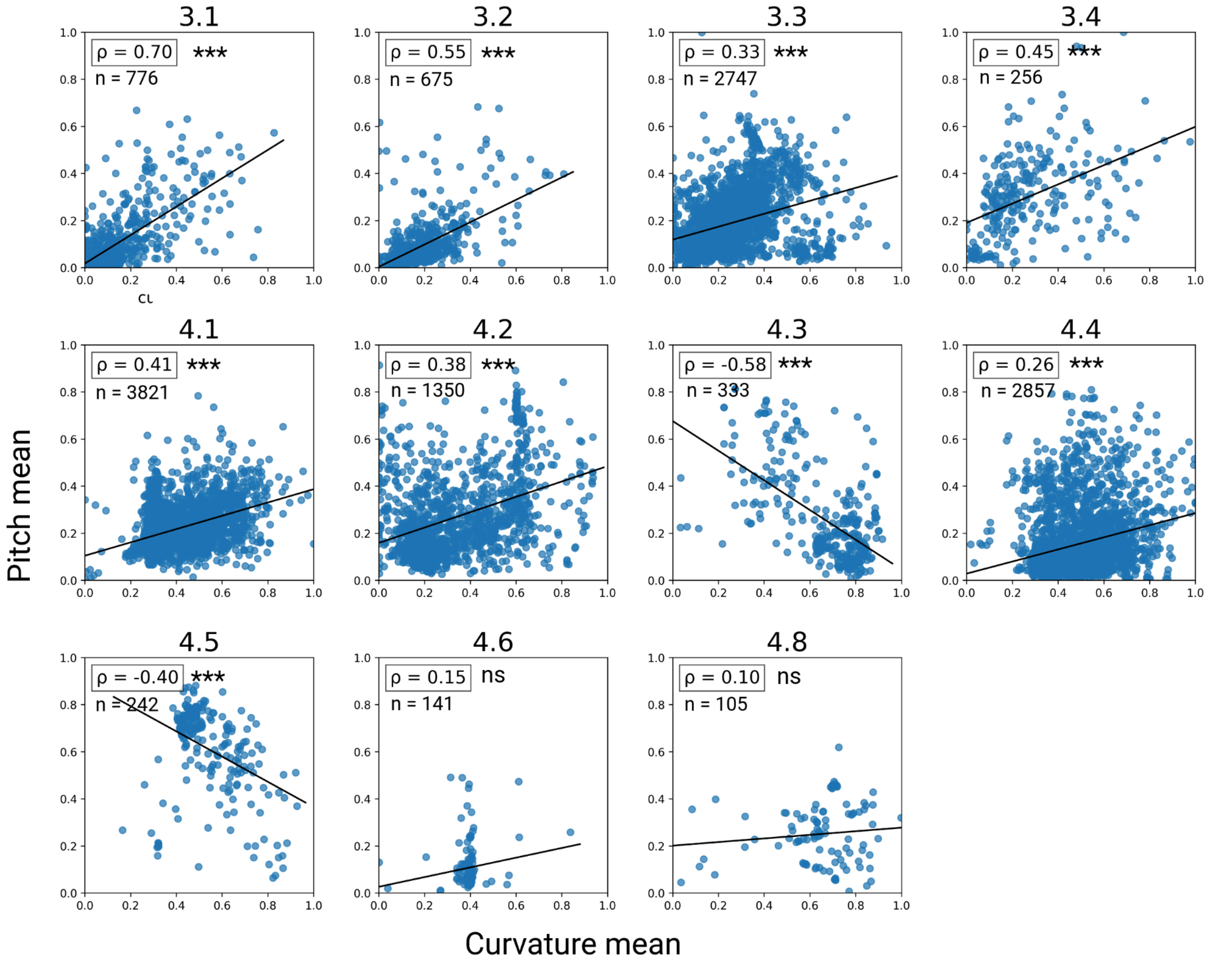


Figure S7. Correlation of mean of curvature and twist parameters for elongated (class 3) and closed (class 4) STRPs at the topology level.

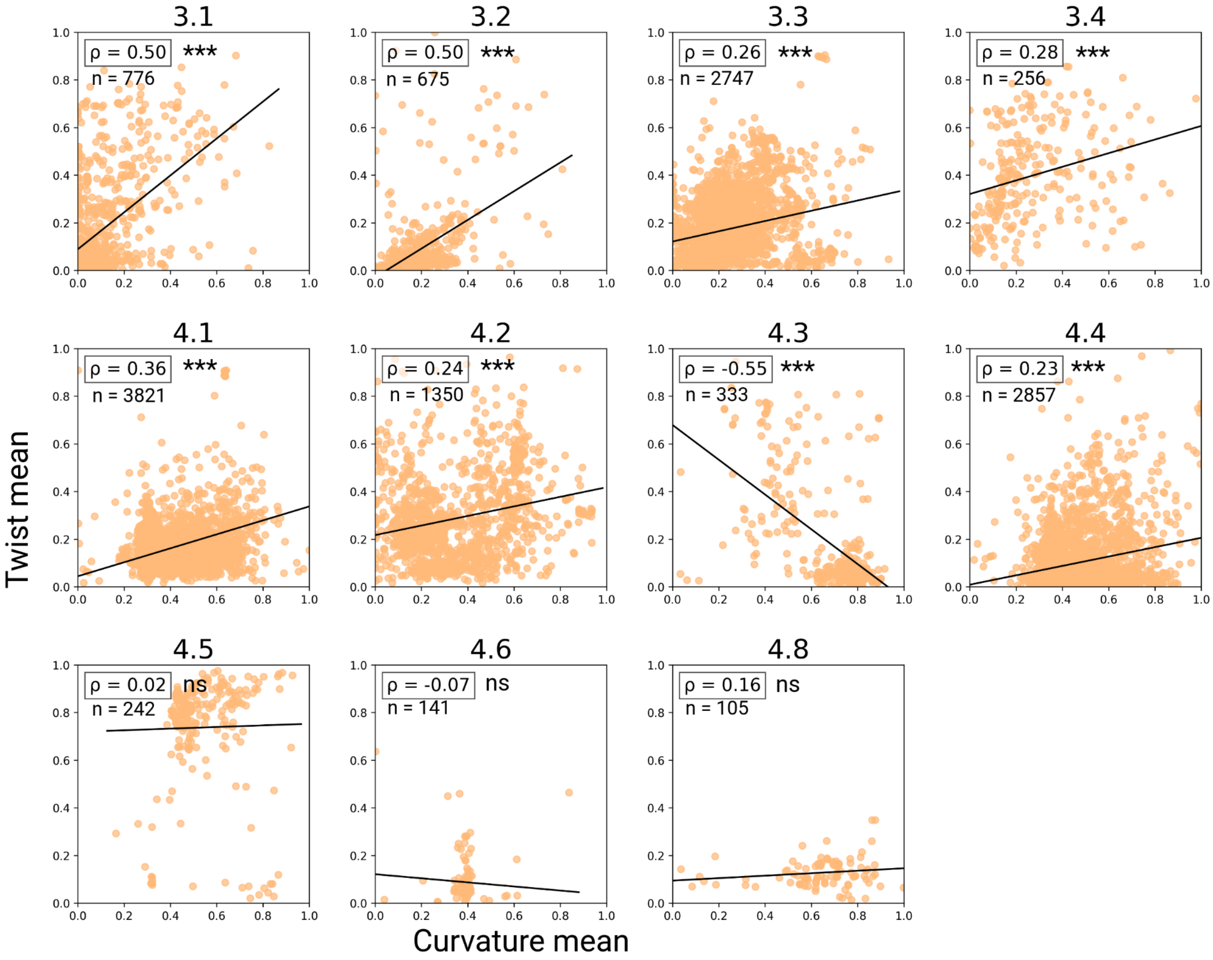


Figure S8. Correlation of mean of curvature and twist parameters for elongated (class 3) and closed (class 4) STRPs at the topology level.

### 4. Mean and standard deviation (std) correlations

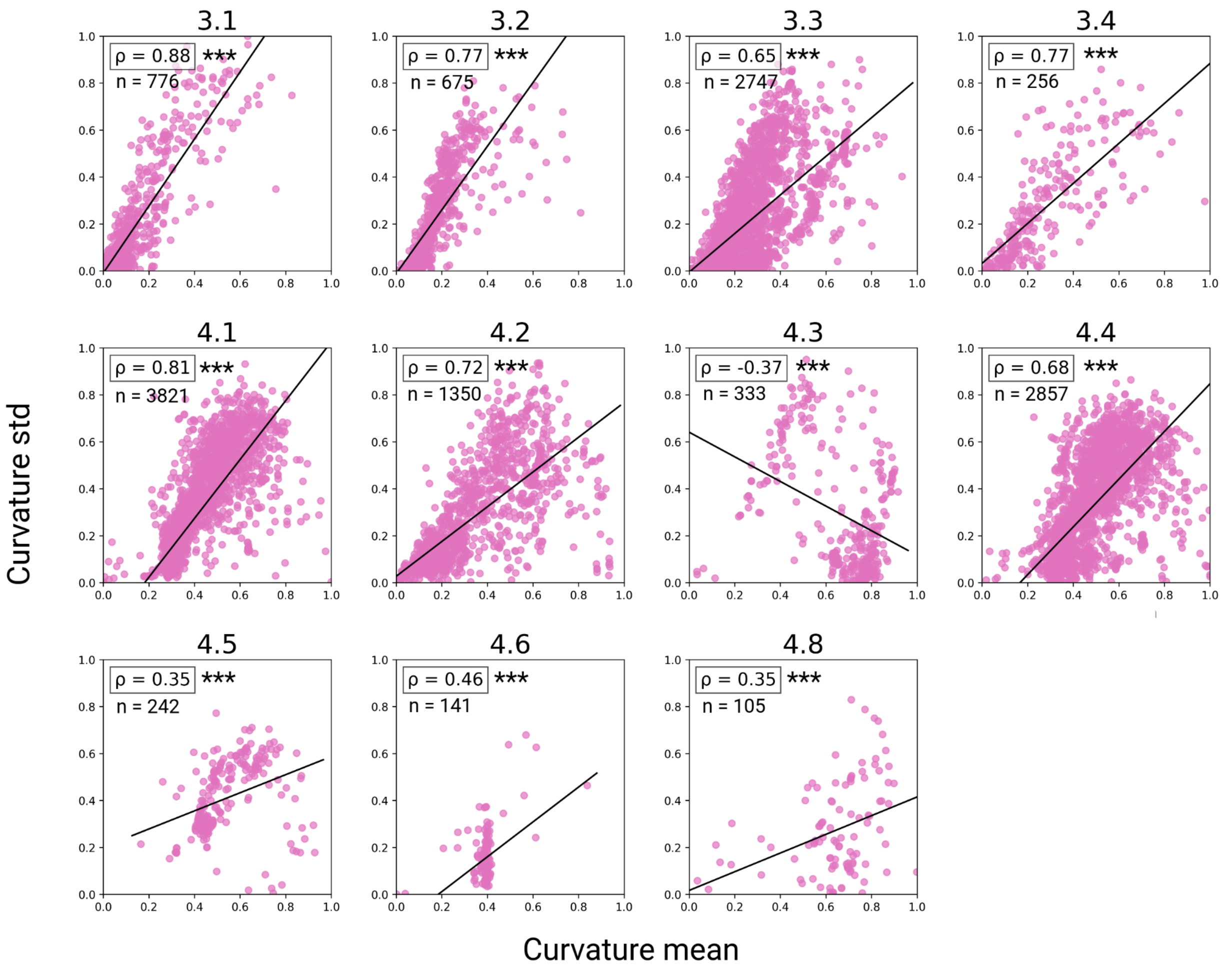


Figure S9. Correlation distribution of mean and standard deviation of curvature across topologies of elongated (class 3) and closed (class 4) STRPs.

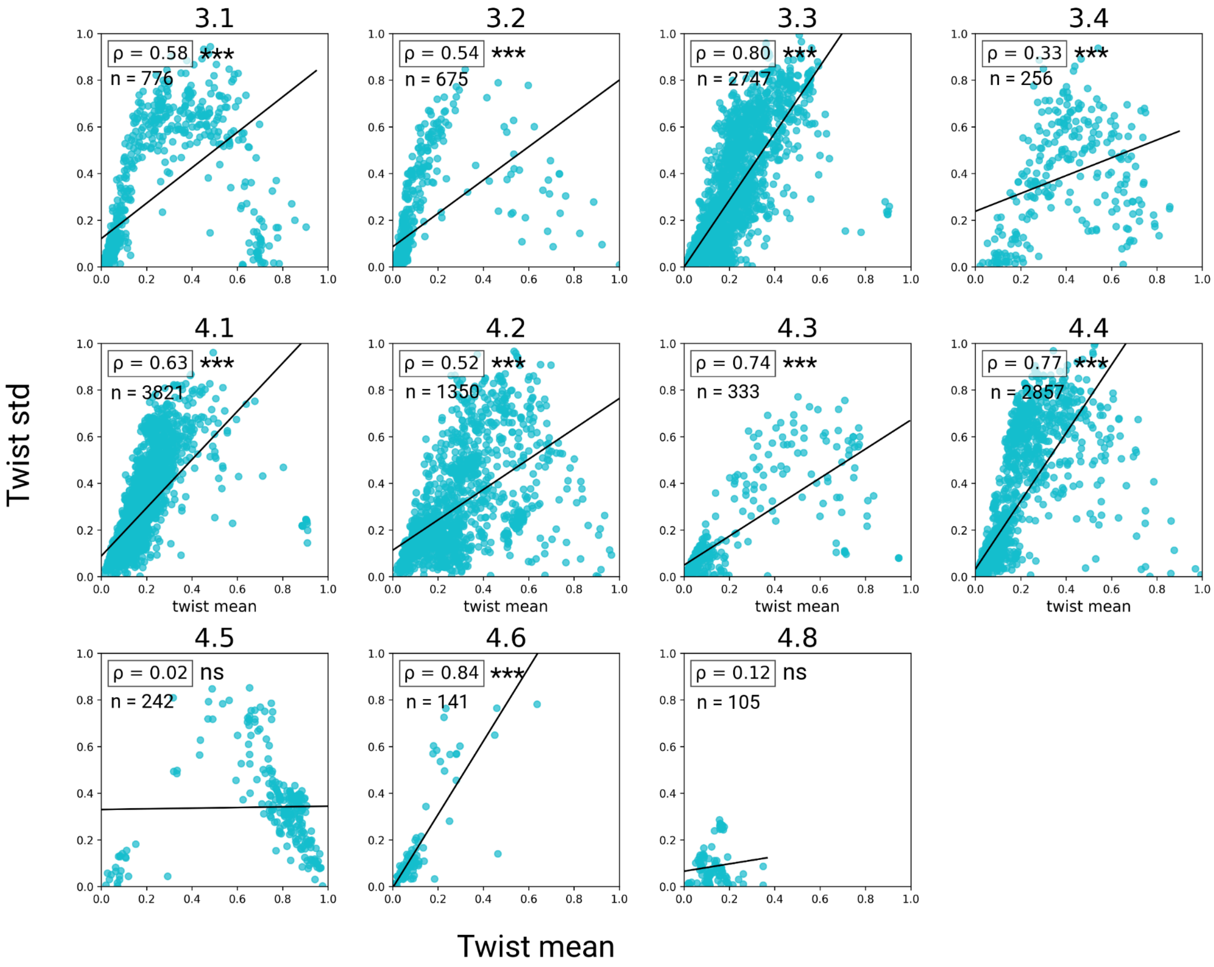


Figure S10. Correlation distribution of mean and standard deviation of twist across topologies of elongated (class 3) and closed (class 4) STRPs.

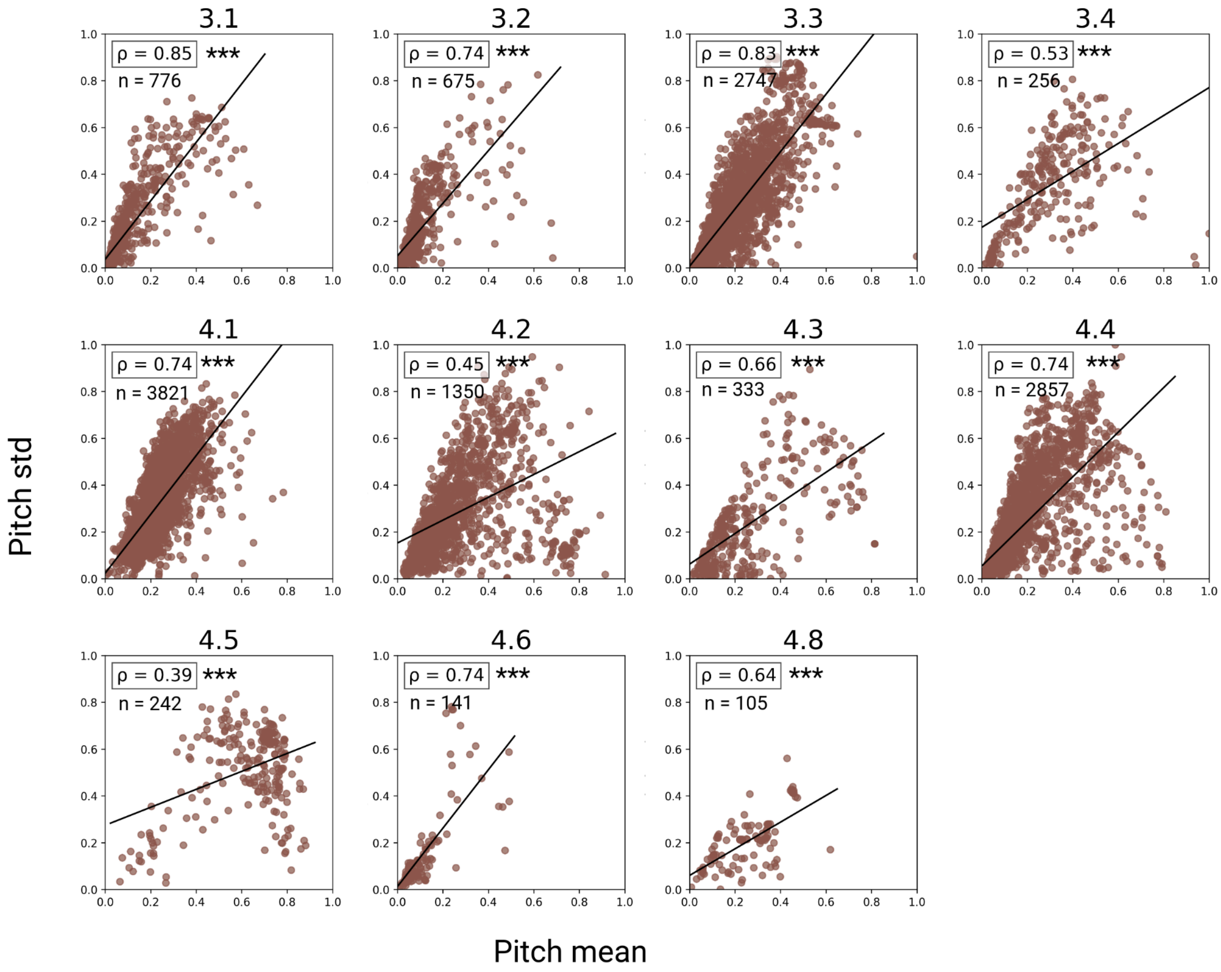


Figure S11. Correlation distribution of mean and standard deviation of pitch across topologies of elongated (class 3) and closed (class 4) STRPs.

### 5. Comparison between natural and designed STRPs

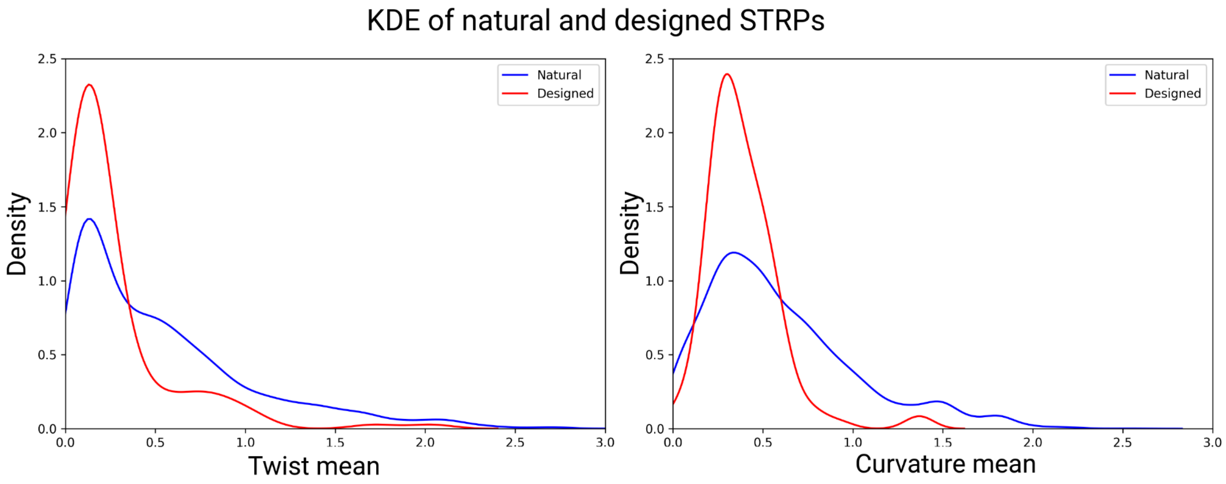


Figure S12. Kernel Density Estimate (KDE) distribution of twist and curvature parameters for natural and designed STRPs of class 3 and class 4.

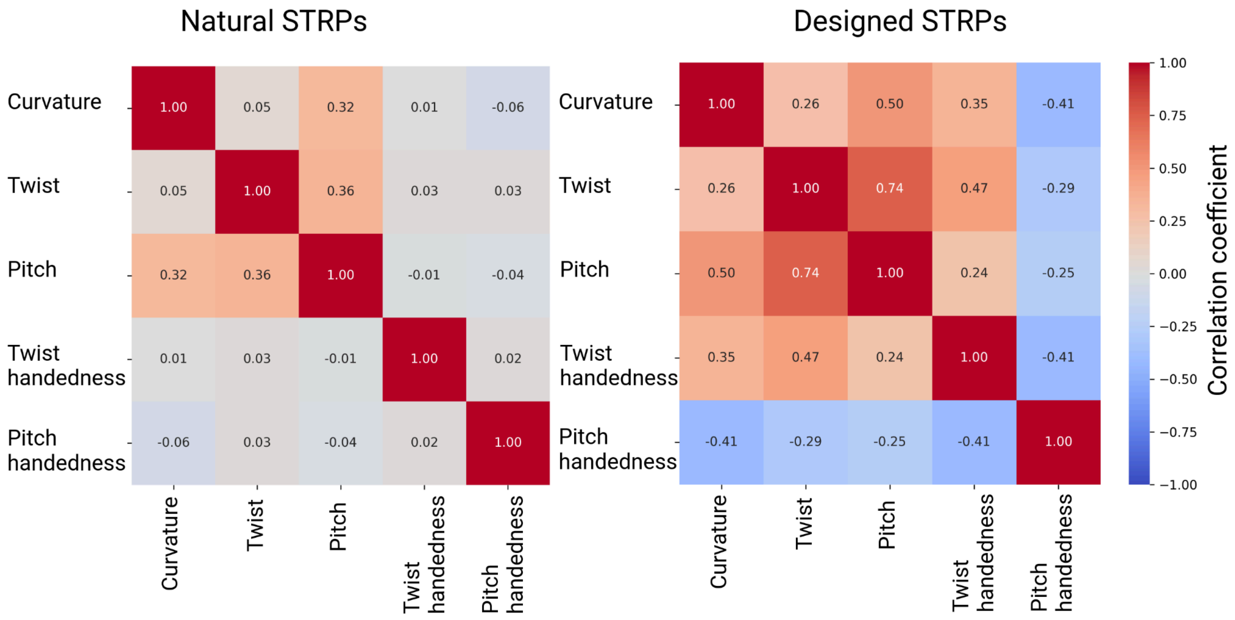


Figure S13. Heatmap of Pearson Correlation of geometry parameters (curvature, twist, pitch, twist handedness, pitch handedness) in natural and designed STRPs.
